# Top-down expectation effects of food labels on motivation

**DOI:** 10.1101/199265

**Authors:** Joost Wegman, Ilke van Loon, Paul A.M. Smeets, Roshan Cools, Esther Aarts

**Affiliations:** Radboud University, Donders Institute for Brain, Cognition and Behaviour, Centre for Cognitive Neuroimaging (Nijmegen, Netherlands); Wageningen University and Research, Division of Human Nutrition (Wageningen, Netherlands); University Medical Center Utrecht, Image Sciences Institute (Utrecht, Netherlands)

## Abstract

When we buy our food, the information on the package informs us about the properties of the product, such as its taste and healthiness. These beliefs can influence the processing of food rewards and impact decision making beyond objective sensory properties. However, no studies, within or beyond the food domain, have assessed how written information, such as food labels, affect implicit motivation to obtain rewards, even though choices in daily life might be strongly driven by implicit motivational biases. We investigated how written information affects implicit motivation to obtain food rewards. We used food labels (high- and low-calorie), associated with an identical lemonade, to study motivation for food rewards during fMRI. In a joystick task, hungry participants (N=31) were instructed to make fast approach or avoid movements to earn the cued drinks. Behaviorally, we found a general approach bias, which was stronger for the drink that was most preferred during a subsequent choice test, i.e. the one labeled as low-calorie. This behavioral effect was accompanied by increased BOLD signal in the sensorimotor cortex during the response phase of the task for the preferred, low-calorie drink compared with the non-preferred, high-calorie drink. During the anticipation phase, the non-preferred, high-calorie drink label elicited stronger fMRI signal in the right ventral anterior insula, a region associated with aversion and taste intensity, than the preferred, low-calorie label. Together, these data suggest that high-calorie labeling can increase avoidance of drinks and reduce neural activity in brain regions associated with motor control. In conclusion, we show effects of food labeling on fMRI responses during anticipation and subsequent motivated action and on behavior, in the absence of objective taste differences, demonstrating the influence of written information on implicit biases. These findings contribute to our understanding of implicit biases in real-life eating behavior.

## Introduction

We know how to behave not only through experience, but also through top-down information, such as instructions. Instructions can control behavior by altering the value of actions, even when such behavior leads to sub-optimal outcomes. For example, misleading information, such as product claims made in marketing, can influence reward-based learning and decision making - and associated neural processes - beyond experience or actual value (Biele et al., 2009; Doll et al., 2011; 2009; Engelmann et al., 2009; Hayes, 1989; Li et al., 2011; Nickerson, 1998). Similarly, misleading information can strongly influence the processing of food rewards. Previous studies have found influences of semantic framing of foods (de Araujo et al., 2005), price labels (Plassmann et al., 2008), healthiness of foods (Grabenhorst et al., 2013; Hare et al., 2011; van der Laan et al., 2012), and "organic" labels (Linder et al., 2010) on neural processing of passively tasted foods or on explicit choice and valuation measures such as liking and willingness-to-pay. Interestingly, the top-down influences of food labels impacts explicit motivation in children (Enax et al., 2015) and can even act on low levels in the gut-brain axis, such as secretion of the gut peptide ghrelin, which signals hunger state to the brain (Crum et al., 2011). However, no studies within or beyond the food domain, have assessed how written information, such as food labels, affect the implicit motivation to obtain rewards, as well as underlying neural processes. This is relevant as behavior is motivated not only by explicit goals that people set for themselves, but also by implicit biases that do not necessarily contribute to adaptive, optimized behavior. Approach biases, defined as an automatic behavioral inclination to approach rather than avoid certain stimuli, can lead to drug-seeking behavior in addiction, despite negative consequences (Watson et al., 2012). Here, we assess the degree to which written information affects approach bias towards food rewards in healthy participants.

Approach biases have been demonstrated by having participants make approach versus avoid movements with a joystick upon stimulus presentation, such as pictures (Rinck and Becker, 2007; Watson et al., 2012). This paradigm has also been successfully applied to study approach biases in the food domain. Obese relative to lean subjects showed enhanced approach biases towards food cues (Havermans et al., 2011; Kemps and Tiggemann, 2015). Furthermore, approach biases to food stimuli were observed when people were food deprived (Seibt et al., 2006), and when food stimuli were appealing rather than disgusting (Piqueras-Fiszman et al., 2014). However, whether behavioral approach biases are affected by written labels is unclear, as well as the neural mechanisms underlying this effect.

Specifically, it is unknown whether the neural counterpart of top-down label effects on approach-avoidance can be expected in regions involved in taste processing (Woods et al., 2011), motivational processing (Cousijn et al., 2012; Doll et al., 2009) or motor control (Mogenson et al., 1980; Radke et al., 2016; Salamone and Correa, 2012). To investigate which brain areas are involved in the effect of written information on the actual motivation to obtain rewards, the present fMRI study examined how approach biases are affected by beliefs. These beliefs were induced by cueing an identical drink as ‘low-calorie’ or ‘high-calorie’, as perceived healthiness of food exerts a strong influence on behavior (Chandon and Wansink, 2012). We employed an approach-avoidance task in which hungry participants worked to actually obtain these differently labeled drinks during fMRI, by responding to label-independent approach and avoid instructions. Because participants responded in the motivational context of action-dependent outcomes that were given shortly after working for them within the scanner, this task has a high degree of ecological validity. First, we tested whether motivated behavior was influenced by the presented drink label. We expected labeling to modulate approach-versus avoid-related behavior, showing a stronger approach bias for the label people preferred. Next, we assessed neural responses at the moment people worked for the differently labeled drinks and during anticipation based on beliefs about the food, i.e. when presented with the different written labels.

## Methods

### Participants

Our sample consisted of 31 right handed, neurologically and psychologically healthy participants (15 men, mean age = 24, age range = 20-32, mean BMI = 23.1, BMI range 20.3-28.1). Exclusion criteria were: a BMI <18.5 or >30 kg/m2, problems with chewing or swallowing, stomach or bowel diseases, diabetes, thyroid disease or any other endocrine disorder, gaining or losing more than 5kg during the last six months, an energy restricting diet during the past 6 months, having a current alcohol consumption of >28 units per week, being allergic and/or intolerant for products under study or having any contraindication for MRI scanning. We invited 34 participants; data of three participants were discarded because of technical problems. Participants were compensated for participation, and gave written informed consent in a manner approved by the local ethics committee on research involving human participants (CMO region Arnhem-Nijmegen, The Netherlands).

### Experimental procedure

Participants were instructed to fast for at least 6h prior to the experiment (no food, only water). Participants were told that they would be working to earn different drink rewards by making correct and fast responses on a joystick task. Inside the scanner, the experiment started with a training session in which participants were familiarised with the experiment and learned the label-taste pairings. This session started with three blocks of 24 trials in which the response requirements of the task (see below) were practiced. Then, the labels were paired with tastes. The labels ‘low-calorie’, ‘neutral’ and ‘high-calorie’ were presented in that order, followed by the presentation of a taste. The neutral label was paired with demineralized water. Unbeknownst to the participants, both the ‘low-calorie’ and ‘high-calorie’ labels were paired with the same lemonade (Karvan Cévitam grenadine, 120g syrup dissolved in 700g of demineralized water; the mixture contained 385 kcal/l). This allowed us to investigate the effects of labelling in the absence of objective taste differences. Each label with the paired drink was delivered three times in 1.5 mL (duration: 3s) quantities, together with a picture indicating the receipt of that drink (a blue drop for the neutral drink, a lighter red for the drink labeled low-calorie and a darker red for the drink labeled high-calorie).

Throughout the experiment, drinks were administered with the use of three identical membrane-liquid pumps (KNF Stepdos FEM03.18RC, KNF Verder, Vleuten, The Netherlands; 0.030-30.0 mL/min) into the participant mouth at a rate of 30 mL/min. After receiving the drinks, participants indicated their liking ("How pleasant do you find the taste of this drink?") and wanting ("How much would you like to drink this drink at this moment?") for the drinks on a continuous VAS scale (ranging from a score of 0 -"Very unpleasant"/"Not at all"- to 10 -"Very pleasant"/"Very much"). Subsequently, a final practice block of 12 trials similar to the actual task (see below) was performed.

Participants were scanned while performing an instrumental approach-avoidance task in which they worked to obtain food rewards (Figure 1). In order to earn the reward cued by the label, participants have to follow the response instructions, i.e. approach (pull) or avoid (push). Because our participants were hungry, we expected them to show an implicit approach bias towards appetitive food stimuli. Trials started with a reward cue (i.e. one of the labels: neutral/low-calorie/high-calorie), which predicted the drink reward for correct performance. The interval between the reward cue and the response target was jittered with a variable delay between 2 and 6 seconds. The target to which participants responded was one of four shapes. For two of the shapes, they had to pull the joystick towards their body (approach), for the other two shapes they had to push the joystick away from their body (avoid). These instructions, with emphasis on the responses relative to their body, were given to them before the start of the scan session and were repeated when they received the joystick while lying on the scanner bed before the start of the experiment.During the experiment, joystick displacements of 80% of the maximal displacement achievable along the sagittal plane were counted as valid responses. To enhance automatic tendencies, responses had to be made within a response deadline, which was adaptive during the experiment: a correct response resulted in a lowering of the deadline with 20 ms, failure to respond within the deadline resulted in an increment of 25 ms. Separate deadlines for each drink cue and each response (approach/avoid) were used. The response deadlines from the end of the practice trials were used as the initial deadlines for the actual experiment. At the time of the response deadline, participants received feedback (“correct”, “incorrect” or “too late”). Feedback “correct” was accompanied by a picture indicating the earned reward. With each correct response, participants earned 0.5 mL of the cued drink.Rewards were not received immediately, but participants received the total amount of liquid they had earned for each drink during each experimental block of 24 trials at the end of that block. This was done to avoid sensory-specific satiety (Rolls et al. 1981), which occurs faster when the same amount of food is received in smaller portions (Weijzen et al. 2009). First, a message was presented that indicated that participants would receive the drinks, then the drink rewards were given in the order: low-calorie, neutral and high-calorie. A vertical bar that decreased in size with drink exposure indicated how much of the drink was still to come. All participants were instructed to refrain from swallowing until instructed to swallow on the screen. After the earned amounts of drinks were presented,0.75 mL of demineralized water was given to rinse. In total, 8 experimental blocks of 24 trials each were presented, making a total of 32 trials per combination of reward cue and response requirement (approach/avoid). After leaving the scanner and performing an unrelated task for about an hour, participants had to choose one of two 0.5 l bottles to take home. The bottles were filled with the lemonade they received during the experiment and looked identical, except for their labels: “‘low-calorie’” and “high-calorie”.

**Figure 1.**
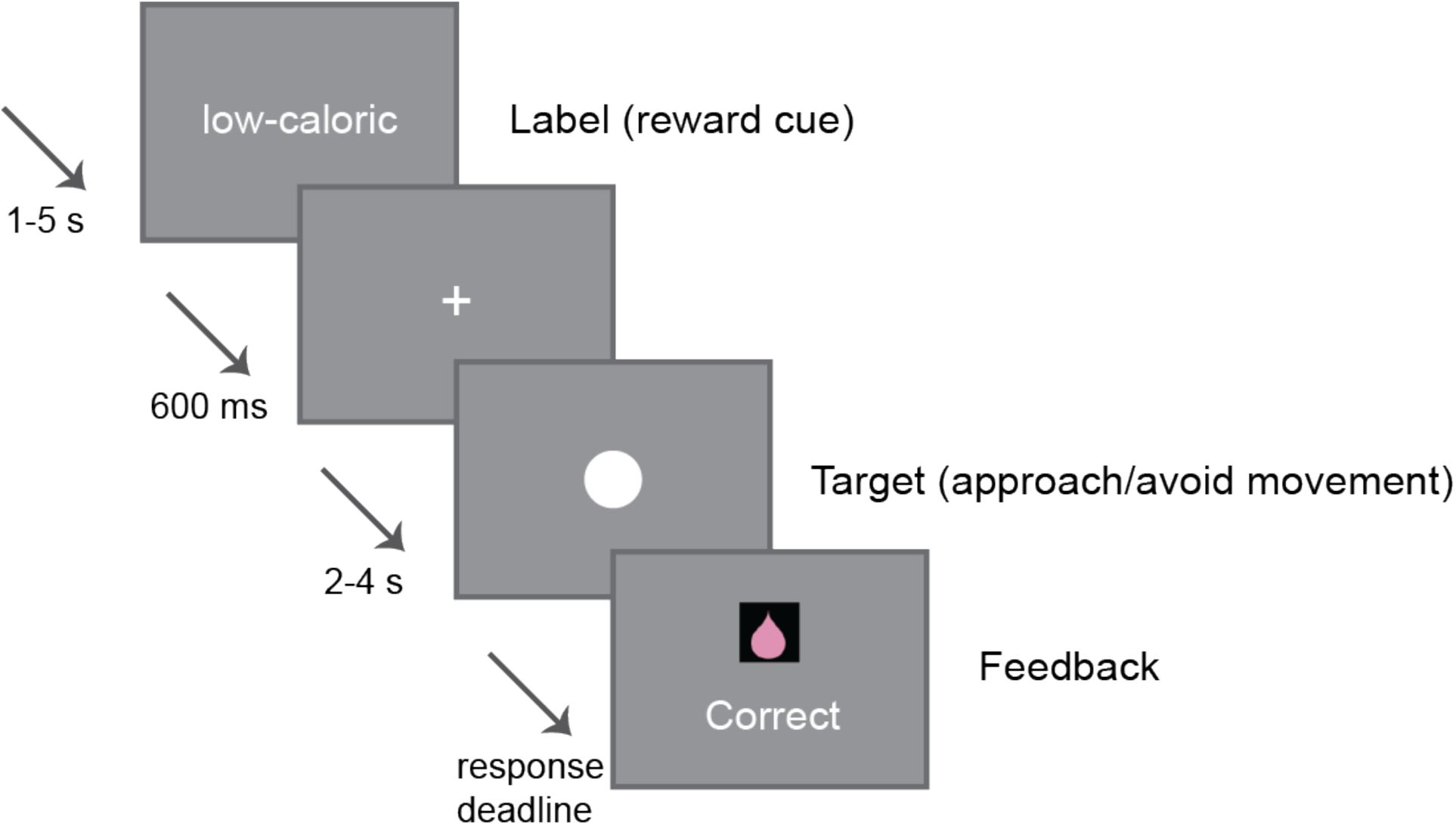
Example trial in the approach bias paradigm. The presentation of one of the reward cue labels (neutral/low-calorie/high-calorie) indicated which reward could be earned that trial. After a variable delay, the response target appeared on the screen. For each shape, participants were trained to make either an approach movement (pull joystick towards their body) or a avoid movement (push joystick away from their body). After the response deadline, feedback was given (correct/incorrect/too late). Rewards were only received at the end of each experimental block.

### Behavioral data analysis

Behavioral data were analyzed using Matlab 8.4 (Mathworks, Natick, MA) and IBM SPSS Statistics (Version 21; IBM, Armonk, NY). To obtain a precise measure of the movement onset (reaction time; RT), joystick movements were reconstructed for each trial using the joystick displacement measurements. Action onsets (RTs) were constrained by several criteria: the joystick needed to be close to the movement onset position (at least < 25% of the maximum deviation [2 cm]) for at least 100 ms prior to onset time, and, subsequently, a sustained deviation for at least 150 ms had to be made. Furthermore, the velocity needed to be significant and sustained (> 0.01 cm/s and peaking to 5 cm/s in the following 100 ms). When these constraints were met, the movement onset was defined as the time-point with the lowest velocity. Criteria for movement offset were defined as: the joystick being close to the maximum offset position (> 80% of the maximum deviation), and at the end of a ballistic movement showing a decreasing velocity (> 5 cm/s in a time window 50-20 ms prior to offset time). Movement time (MT) was defined as the time difference between movement onset and movement offset. We excluded trials that showed extreme RTs (<150ms or >1000ms), RTs and MTs >3 SD from the mean, and trials in which no response was made. For RT analysis, trials in which a response in the wrong direction was made were also excluded. RTs and error rates were analyzed using a repeated-measures general linear model (GLM) with the within-subject factors Action (approach, avoid) and anticipated Labeled drink. Given our experimental aim, we focus on the main contrast of interest between low-calorie and high-calorie labels. For comparison, we also performed analyses that included the neutral label.

## FMRI data analysis

### Data acquisition

Whole-brain functional images were acquired on a 3T Siemens Skyra MRI scanner (Siemens Medical system, Erlangen, Germany) using a 32-channel coil. A multi-echo echo-planar imaging (EPI) sequence was used to acquire 34 axial slices per functional volume (voxel size 3.5 x 3.5 x 3 mm; repetition time 2070 ms; echo times: 9 ms, 19.25 ms, 29.5 ms, and 39.75 ms; flip angle = 90°; field of view = 224 mm). This type of multi-echo acquisition sequence for functional images reduces motion and susceptibility artifacts (Poser et al., 2006). After the acquisition of functional images, a high-resolution anatomical scan was acquired (T1-weighted MPRAGE, voxel size 1 x 1 x 1 mm, TR 2300 ms, TE 3.03 ms, 192 sagittal slices, 1 mm thick, FoV 256 mm).

### Image processing

Data were analyzed using SPM8 (www.fil.ion.ucl.ac.uk/spm). The volumes for each echo time were realigned to correct for motion artifacts (estimation of the realignment parameters was done for the first echo and then copied to the other echoes). The four echo images were combined into a single MR volume based on 30 volumes acquired before the actual experiment started using an optimized echo weighting method (Poser et al., 2006). Combined functional images were slice-time corrected. Structural and functional data were then co-registered and spatially normalised to standard stereotactic space (Montreal Neurological Institute (MNI) space). After segmentation of the structural images, using a unified segmentation approach, the mean of the functional images was spatially coregistered to the bias-corrected structural images. The transformation matrix resulting from segmentation was then used to normalize the final functional images into MNI space (resampled at voxel size 2 x 2 x 2 mm). Finally, the normalized functional images were spatially smoothed using an isotropic 8 mm full-width at half-maximum Gaussian kernel.

### GLM analyses

Statistical analyzes were performed in the context of the general linear model in SPM8. Our first level model contained 9 regressors of interest: 3 regressors for the reward cue labels (neutral, low-calorie, high-calorie) and 6 regressors for the response cues after which a correct response was made (Labeled drink*Action). All regressors of interest were modeled as an impulse response function (duration = 0) convolved with a canonical hemodynamic response function. Furthermore, experiment-related regressors of no interest were included for the response cues to which an incorrect response was made (collapsed over preceding labels), the outcomes (receipt of each type of drink), VAS scales and for rinse and swallow instructions. In addition, we included six motion parameters, their first-order derivatives and global signal changes (as indexed by segmented white matter, cerebral spinal fluid and out-of-brain voxels). Statistical analysis included high-pass filtering (cutoff: 128 seconds) to remove low-frequency confounds such as scanner drifts and correction for serial correlations using an autoregressive AR(1) model.

Contrast images from the first level were entered into second level random-effects analyses to test for consistent effects over participants. We performed different one-sample T-tests based on the following contrasts calculated on the first level: low-calorie vs. high-calorie labels during anticipation. To assess signal during motivated action (approach versus avoid), we calculated the contrast between all approach (pull) instructions > all avoid (push) instructions. Furthermore, to investigate whether implicit biases were affected by the misleading information, we tested for interactions between Labeled drink and Action using the contrast images for the approach bias: (approach vs. avoid) in low-vs. high-calorie labeled trials. The results of all random effects fMRI analyses were thresholded at P< 0.001 (uncorrected) and statistical inference was performed at the cluster level, family-wise-error-corrected (PFWE<0.05) for multiple comparisons over the search volume (the whole brain).

## Results

### Subjective measures

We assessed subjective liking and wanting of the differently labeled drinks using visual analog scales. Liking scores for all three received drinks did not differ at first exposure (F(2,29) = 1.696, P = .201) or when comparing initial tasting with last receipt (main effect of Labeled drink or Labeled drink*Time interaction: Ps >.4). Wanting ratings for the three received drinks also did not differ after the first exposure (F 29,2) = .037, P = .964) or when comparing initial tasting with last receipt (main effect of Labeled drink or Labeled drink*Time interaction: Ps > .35). Overall wanting for drinks decreased over the course of the experiment (main effect of Time: F(1,30)=8.929, P = .006). Excluding the neutral drink from the analysis of liking and wanting ratings yielded the same pattern (i.e. no significant main effects of Labeled drink or Labeled drink*Time interactions: all P > .1).

At the end of the experiment, participants were presented with the choice to take home a 0.5 liter drink bottle with identical lemonade content, labeled as either the low-calorie or the high-calorie drink. Of the 30 participants presented with this choice (for one participant, time constraints prevented this), twenty-three (76.7%) participants chose the bottle with the low-calorie label (χ^2^(1)=14.226, P<.001). Therefore, we interpret the low-calorie label as being the preferred label.

Finally, to test whether the instructed knowledge manipulation worked, we asked participants to describe the difference between the drinks labeled as low-calorie and those labeled as high-calorie. Of the 30 participants who filled out this question, 26 reported that they had consistently tasted a difference (χ^2^(1)=16.133, P<.001). Of these 26 participants, 22 described the drink labeled high-calorie as being sweeter/more intense, 3 described the drink labeled low-calorie as being sweeter/more intense, and 1 did not specify which one was perceived as such (label described as sweeter: χ^2^(1)=14.440, P<.001).

### Behavioral results

First, we assessed whether error rates and reaction times (RTs) of the approach (i.e. pull) and avoid (i.e. push) joystick actions depended on the preceding reward cues (i.e. neutral, or high or low-calorie label). Overall, participants made fewer errors when instructed to make approach versus avoid actions (main effect of Action F(1,30)=30.0, P<.001).

Importantly, this approach bias depended on the anticipated label (Labeled drink*Action F(2,29)=20.619, P<.001; Table 1). Breakdown of this interaction showed a significant approach bias for the low-calorie (T(30)=5.747, P<.001) and neutral (T(30)=4.013,P<.001) labels and a marginally significant effect for the high-calorie label (T(30)=1.965,P=.059). Simple main effects analyses revealed that, after approach instructions, significantly fewer errors were made on low-than high-calorie trials (T(30)=3.445,P<.001). There was a trend towards fewer errors on neutral than high-calorie trials (T(30)=1.838,P=.076), and no difference between low-calorie and neutral trials (T(30)=.461,P>.6). After avoid instructions, significantly more errors were made on low-calorie than high-calorie trials (T(30)=2.867,P<.01) and on neutral than high-calorie trials (T(30)=2.815,P<.01), whereas no difference was observed between neutral and low-calorie trials (T(30)=.045, P>.9). Thus, the Labeled drink*Action interaction was driven primarily by greater approach bias for the preferred (low-calorie) versus non-preferred (high-calorie) labels (Figure 2). To substantiate the interpretation that the low-calorie label was preferred on the behavioral level, we performed a post-hoc analysis within the subgroup of participants that all chose (i.e. preferred) the drink labeled low-calorie at the end of the experiment (n=23). Within this group, we observed the same Labeled drink*Action interaction over all three Labeled drinks (F(2,21)=9.967,P<.001). Simple main effects in this group confirmed that significantly fewer errors were made on low-(preferred) than high-calorie (non-preferred) trials (T(22)=2.833,P=.01), whereas no difference was observed between neutral and either low or high calorie labels (Ps>.29). After avoid instructions, this subgroup also made significantly more errors on low-calorie (preferred) than high-calorie (non-preferred) trials (T(22)=2.454,P=.022) and on neutral than high-calorie trials (T(22)=3.008,P<.01), whereas no difference was observed between neutral and low-calorie trials (T(22)=.167,P>.8).

**Table 1.**
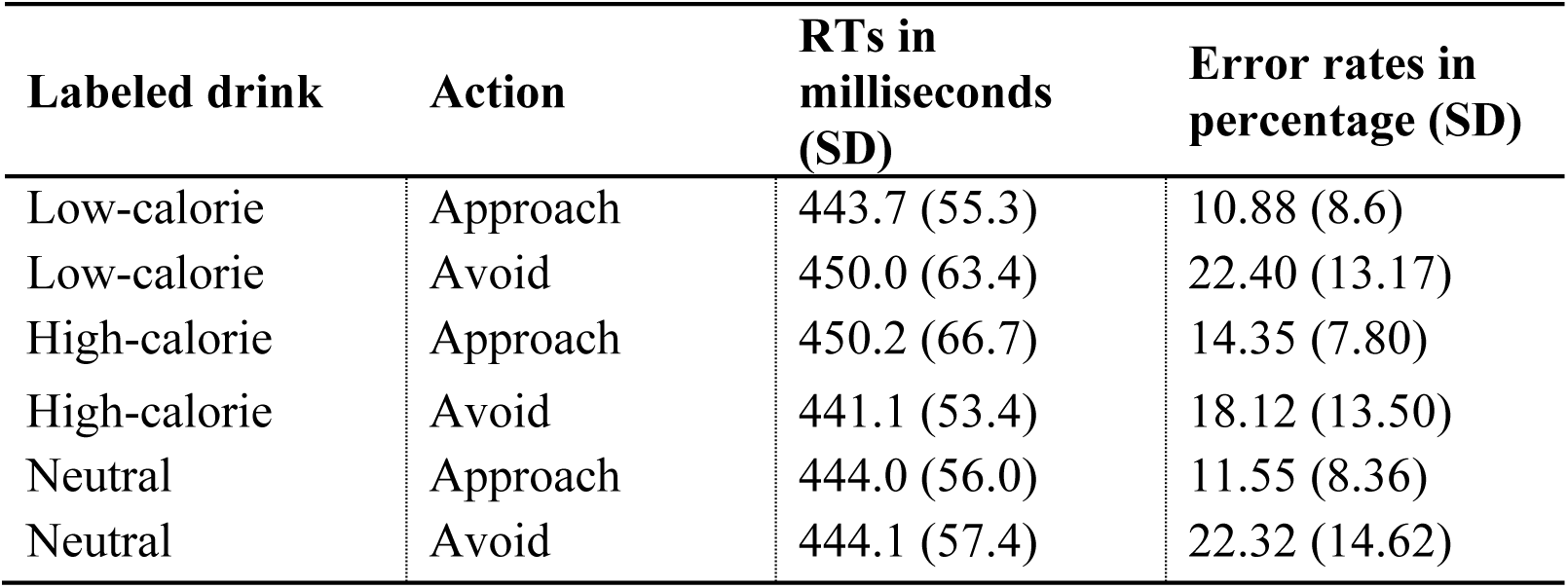
Behavioral results. RTs (in milliseconds) and error rates (in percentages) for each combination of reward cue label and instructed movement.

| Labeled drink | Action | RTs in milliseconds (SD) | Error rates in percentage (SD) |
| --- | --- | --- | --- |
| Low-calorie | Approach | 443.7 (55.3) | 10.88 (8.6) |
| Low-calorie | Avoid | 450.0 (63.4) | 22.40 (13.17) |
| High-calorie | Approach | 450.2 (66.7) | 14.35 (7.80) |
| High-calorie | Avoid | 441.1 (53.4) | 18.12 (13.50) |
| Neutral | Approach | 444.0 (56.0) | 11.55 (8.36) |
| Neutral | Avoid | 444.1 (57.4) | 22.32 (14.62) |

**Figure 2.**
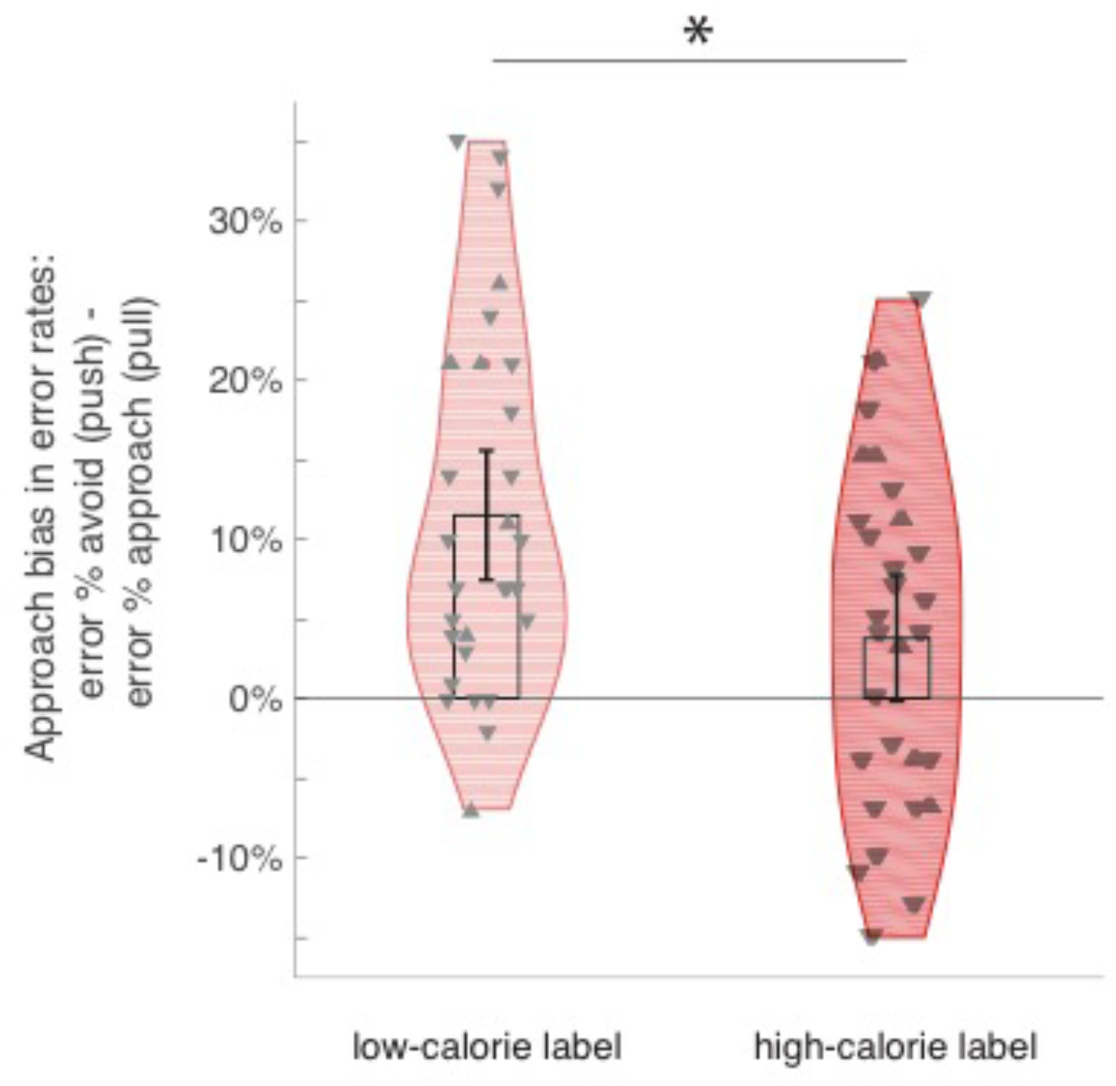
Approach bias in error rates: error percentages for avoid (push) minus approach (pull) instructions, made during working for the low-calorie and high-calorie labeled drinks. Markers indicate whether participants chose the drink labeled low-calorie (▼) or high-calorie (▲) at the end of the experiment (see Materials and Methods).

An ANOVA of RTs to all drink labels did not show any Labeled drink*Action interaction (F(2,29)=2.08, P=.143), nor any main effect (both Fs <1). Given our main contrast of interest was between the calorie labels, we proceeded without the neutral label. In this statistical model, RTs for approach versus avoid actions differed depending on the cued reward label (Labeled drink*Action: F(1,30)=4.287, P=.047; see Table 1). This was due to slower avoid actions after a low-versus high-calorie label (T(30)=2.079, P=.046), but no difference in RTs for approach actions after low-versus high-calorie labels (T(30)=-1.103, P=.279). No main effects of Labeled drink or Action were observed for the RTs (F(1,30)<1). We again performed a post-hoc analysis within the subgroup of participants that all chose (i.e. preferred) the drink labeled low-calorie at the end of the experiment (n=23) to support the interpretation that the low-calorie label was preferred on the behavioral level. Again, we did not see any Labeled drink*Action interaction (F(2,21)=1.571, P=.231), nor any main effect (both Fs <1). When disregarding the neutral label as before, we find a marginally significant Labeled drink*Action interaction (F(2,21)=3.284,P=.084). This trending effect seems to be driven by a trend towards slower avoid actions after a low-versus high-calorie label (T(22)=1.836, P=.08), whereas no difference in RTs for approach actions after low-versus high-calorie labels was observed (T(22)=-.974, P=.341).

In sum, behavioral approach tendencies and choices depended on the labels of the drink, despite the absence of any objective value differences (i.e. same lemonade). Specifically, we observed a greater approach bias (i.e. more approach than avoid movements despite an equal amount of approach and avoid instructions) for the preferred (low-calorie) than non-preferred (high-calorie) label. This effect was paralleled by slower RTs when having to make an avoid movement for the preferred (low-calorie) than non-preferred (high-calorie) label. These behavioral results are therefore in line with preferences, indicated by choices for labeled drinks made after the experiment. The link with subject preference is supported by subgroup analyses based on these choices.

## fMRI results

### Anticipation effects of label

To identify the brain regions involved in label-related anticipation, we compared BOLD responses to high-calorie labels with that during low-calorie labels. This comparison showed significant clusters in left middle temporal gyrus, left superior temporal pole and a cluster overlapping the right temporal pole and the right ventral anterior insula (Figure 3 and Table 2a). No significant clusters were found for the contrast low-calorie label > high-calorie label. Both the low-calorie > neutral and high-calorie > neutral comparisons yielded responses in occipitotemporal cortex, which could be due to the larger visual input associated with the longer labels (i.e. words). No results were found in other regions or for the reverse contrasts. Given that the insula is involved in taste processing and food rewards (Dalenberg et al., 2015; Sescousse et al., 2013; van der Laan et al., 2011; Woods et al., 2011) but also in processing aversive stimuli (Bermúdez-Rattoni, 2014; Nitschke et al., 2006a; Wicker et al., 2003), we performed a post-hoc contrast on the anticipated labels in the subgroup of participants that all chose (i.e. preferred) the drink labeled low-calorie at the end of the experiment (n=23; cluster-defining threshold P<0.001; k>10). In this subgroup, the contrast high-calorie (non-preferred) label > low-calorie (preferred) label yielded whole-brain significant (pFWE<.05) clusters in left anterior temporal cortex and left medial temporal lobe, overlapping hippocampus and amygdala, as well as (uncorrected; P<.001, k>10) activations in other temporal regions and putamen (Table 2b).

**Figure 3.**
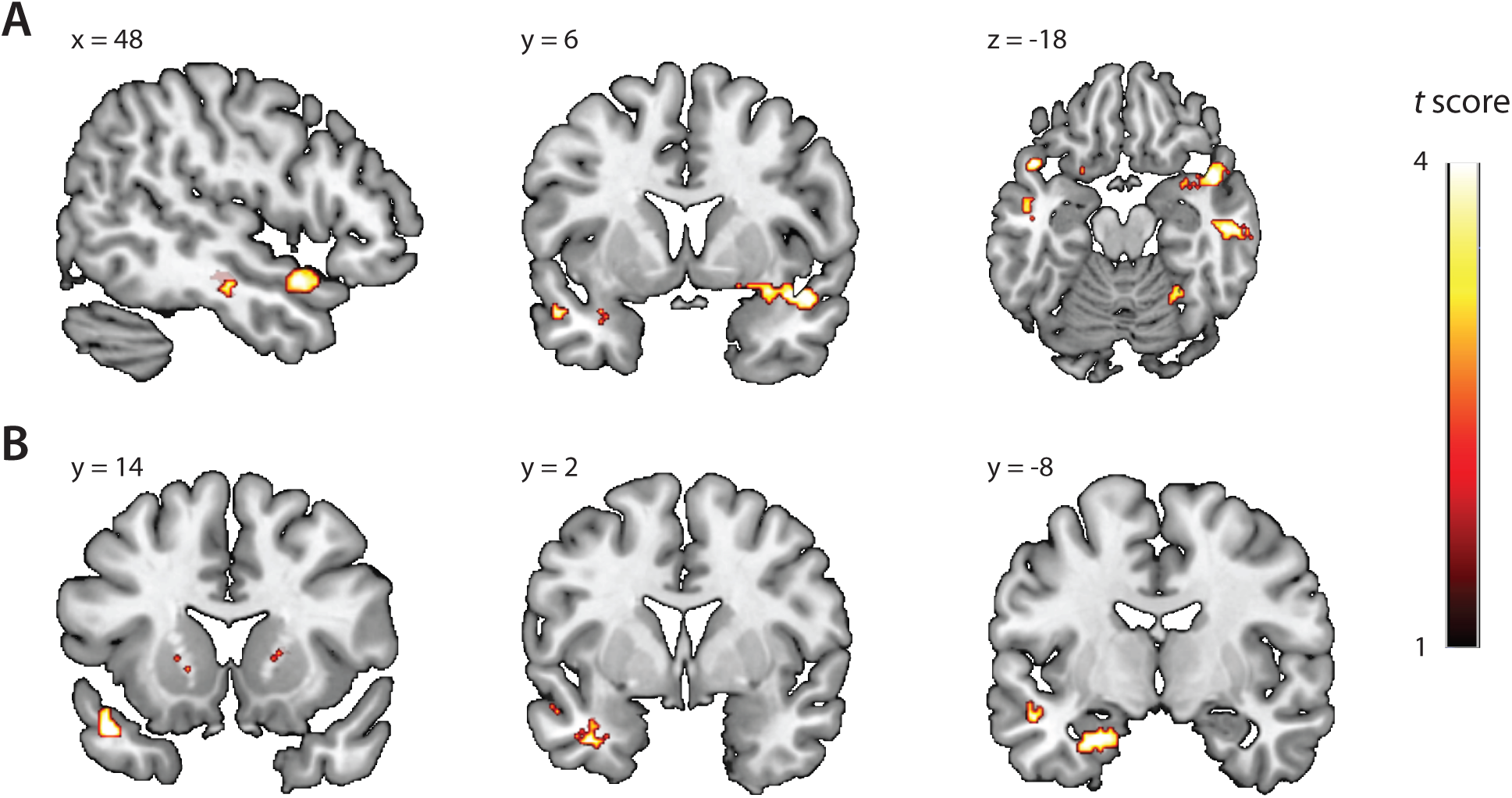
Anticipation-related brain responses as a function of drink label. **A**) High-calorie > low-calorie in the whole sample (N=31). **B**) Non-preferred (high-calorie) > preferred (low-calorie) within the subgroup of participants that all chose (i.e. preferred) the drink labeled low-calorie at the end of the experiment (n=23). Activations shown at P < .001, k>10. For coordinates and which activations survive P_FWE_ < 0.05 at the cluster level (*), see Table 2a and 2b.

**Table 2a.** Brain regions showing significant response during the anticipation phase of trials (*P_FWE_ < 0.05 at the whole-brain corrected cluster level; cluster-defining threshold: P < .001).

| Contrast/Region | Cluster |  |  |  |  |
| --- | --- | --- | --- | --- | --- |
|  | size<br>(voxels) | x | y | z | peak t |
| ***Anticipation high-calorie > low-calorie label*** |  |  |  |  |  |
| L middle temporal gyrus | 146 | -52 | -14 | -10 | 5.40* |
| R superior temporal pole | 164 | 48 | 6 | -18 | 4.60* |
| R ventral insula |  | 38 | 6 | -16 |  |
| R inferior/middle temporal gyrus | 117 | 54 | -20 | -18 | 4.59* |
| ***Anticipation low-calorie > neutral label*** |  |  |  |  |  |
| L/R occipitotemporal cortex | 4793 | -14 | -90 | 0 | 8.29* |
| ***Anticipation high-calorie > neutral label*** |  |  |  |  |  |
| L occipitotemporal cortex | 4547 | -16 | -88 | -10 | 8.84* |
| R occipitotemporal cortex | 755 | 18 | -90 | -4 | 7.97* |

**Table 2b.** Brain regions showing significant response during the anticipation phase of trials within the subgroup of participants that all chose (i.e. preferred) the drink labeled low-calorie at the end of the experiment (n=23; threshold: P < .001, k>10; *P_FWE_ < 0.05 at the whole-brain corrected cluster level).

| **Contrast/Region** | **Cluster** |  |  |  |  |
| --- | --- | --- | --- | --- | --- |
|  | **size**<br>**(voxels)** | **x** | **y** | **z** | **peak t** |
| ***Anticipation high-calorie (non-preferred) > low-calorie (preferred) label in subgroup that chose low-calorie drink (n=23)*** |  |  |  |  |  |
| L anterior temporal lobe | 185 | -46 | 14 | -24 | 5.00* |
| L middle temporal cortex |  | -52 | -14 | -10 |  |
| R fusiform gyrus/hippocampus | 57 | 36 | -28 | -18 | 4.73 |
| R parahippocampal gyrus |  | 26 | -26 | -20 |  |
| L medial temporal lobe/amygdala | 174 | -36 | 2 | 32 | 4.65* |
| L anterior hippocampus/amygdala |  | -30 | -8 | -28 |  |
| L inf temporal cortex |  | -42 | -4 | -32 |  |
| R inf temporal cortex | 51 | 48 | -14 | -24 | 4.31 |
| L putamen | 75 | -22 | 8 | 4 | 4.29 |
| R putamen | 13 | 20 | 16 | 4 | 4.21 |
| R anterior temporal lobe | 19 | 48 | 10 | -20 | 4.05 |
| ***Anticipation low-calorie (preferred) > high-calorie (non-preferred) label in subgroup that chose low-calorie drink (n=23)*** |  |  |  |  |  |
| R frontal white matter | 10 | 26 | 26 | 16 | 4.56 |

### Effects of top-down labeling on neural signaling during motivated action

We first assessed the main effect of motivated action, that is the difference between approach and avoid actions. Signal was reduced during approach versus avoid actions in the cerebellum and a cluster in left sensorimotor cortex (postcentral gyrus), in line with responding with their right hand (Table 3; Figure 4). Next, we assessed the degree to which neural signals during motivated action were modified by top-down drink labeling. The left sensorimotor cortex (in the postcentral gyrus, not overlapping the cluster in left sensorimotor cortex observed in the main avoid > approach contrast), as well as the left superior parietal cortex exhibited a Labeled drink*Action interaction, due to stronger approach versus avoid responses after the low-calorie label versus the high-calorie label (Table 3; Figure 4). For the reverse interaction (approach > avoid after high-calorie > low-calorie label), we observed a cluster in bilateral occipitotemporal cortex. In keeping with the Labeled drink (low vs high) x Action contrast, a comparison between the low-calorie trials with the neutral label trials also revealed a Labeled drink*Action interaction in the postcentral gyrus: approach versus avoid signal in this region was greater for low-calorie label trials than for neutral trials. There were no main effects of Labeled drink irrespective of Action during the response phase. To assess the degree to which these results are driven by the preference of participants, we performed a post-hoc contrast on the Labeled drink*Action interaction in the subgroup of participants that all chose (i.e. ‘preferred’) the drink labeled low-calorie at the end of the experiment (n=23; cluster-defining threshold P<0.001; k>10). In this subgroup, approach > avoid after high-calorie (non-preferred) > low-calorie (preferred) label, we did not find whole-brain significant (pFWE<.05) clusters, but we did find activations in similar sensorimotor regions as those observed in the whole sample uncorrected at the whole-brain level (P<0.001, k>10; Table 3b). For the reverse interaction (approach > avoid after high-calorie > low-calorie label), we observed a range of occipitotemporal visual regions, as we did in the entire sample.

**Figure 4.**
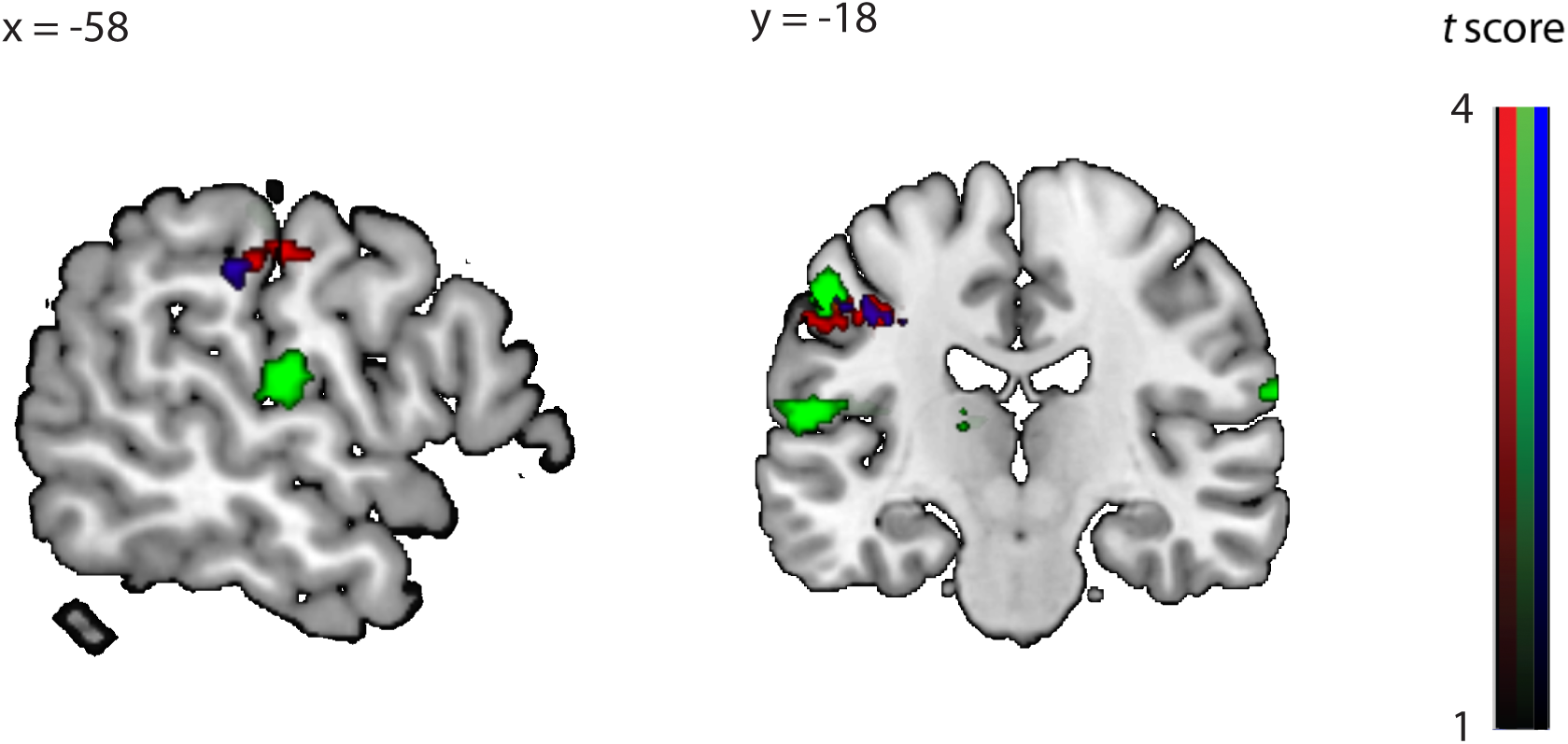
Motivated action-related brain responses. The Labeled drink*Action interaction (approach > avoid: low-calorie > high-calorie) is shown in red (N=31). **T**he contrast of all avoid (push) > approach (pull) actions is shown in green (N=31). The Labeled drink*Action interaction (approach > avoid: preferred (low-calorie) > non-preferred (high-calorie) within the subgroup of participants that all chose (i.e. preferred) the drink labeled low-calorie at the end of the experiment (n=23) is shown in blue. Activations shown at P < .001, k>10. For coordinates and which activations survive P_FWE_ < 0.05 at the cluster level (*), see Table 3a and 3b.

**Table 3a.**
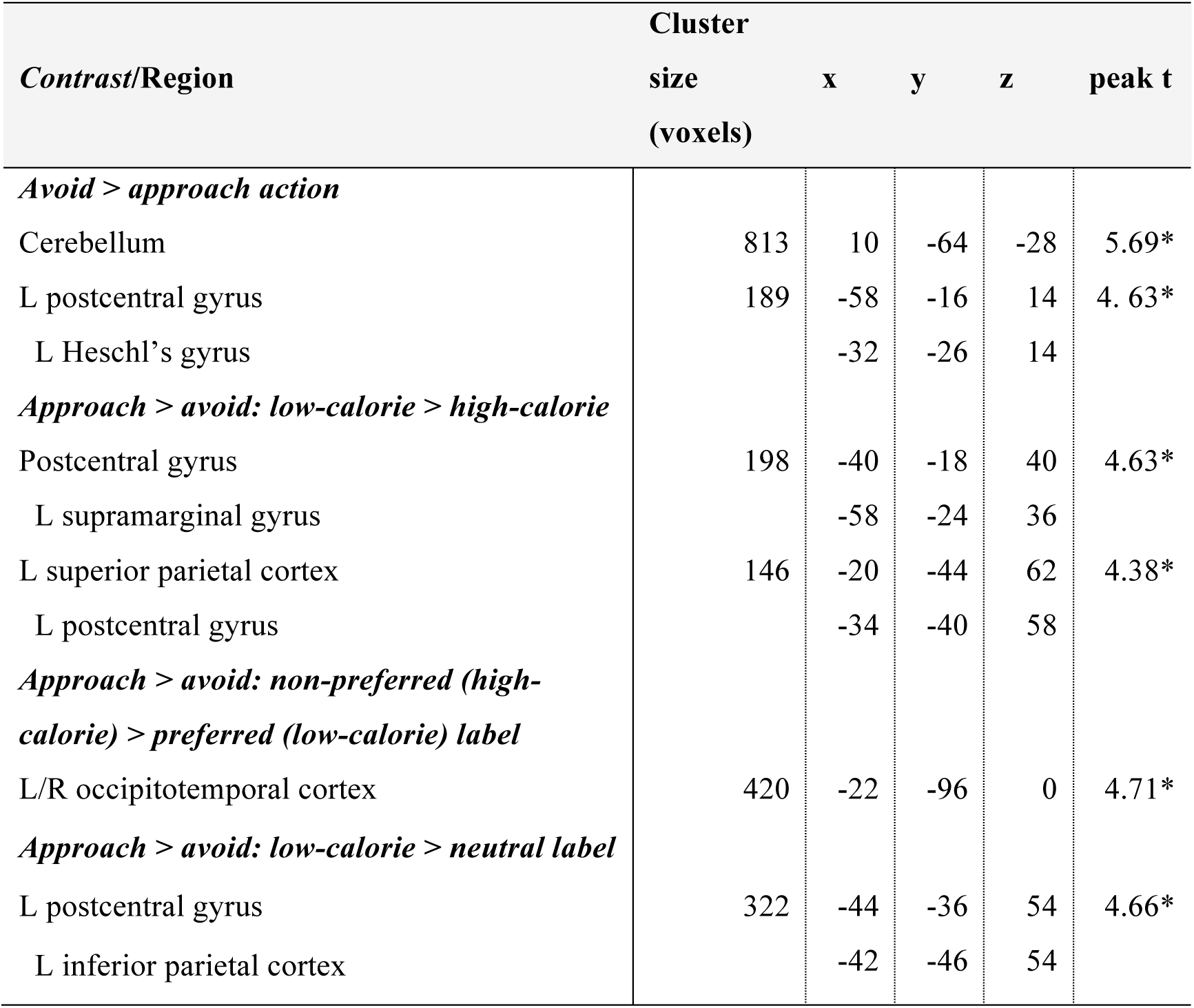
Brain regions showing signification responses during the response phase of trials (*P_FWE_ < 0.05 at the whole-brain corrected cluster level; cluster-defining threshold: P < .001).

| **Contrast/Region** | **Cluster**<br>**size**<br>**(voxels)** | **x** | **y** | **z** | **peak t** |
| --- | --- | --- | --- | --- | --- |
| ***Avoid > approach action*** |  |  |  |  |  |
| Cerebellum | 813 | 10 | -64 | -28 | 5.69* |
| L postcentral gyrus | 189 | -58 | -16 | 14 | 4.63* |
| L Heschl's gyrus |  | -32 | -26 | 14 |  |
| ***Approach > avoid: low-calorie > high-calorie*** |  |  |  |  |  |
| Postcentral gyrus | 198 | -40 | -18 | 40 | 4.63* |
| L supramarginal gyrus |  | -58 | -24 | 36 |  |
| L superior parietal cortex | 146 | -20 | -44 | 62 | 4.38* |
| L postcentral gyrus |  | -34 | -40 | 58 |  |
| ***Approach > avoid: non-preferred (high-calorie) > preferred (low-calorie) label*** |  |  |  |  |  |
| L/R occipitotemporal cortex | 420 | -22 | -96 | 0 | 4.71* |
| ***Approach > avoid: low-calorie > neutral label*** |  |  |  |  |  |
| L postcentral gyrus | 322 | -44 | -36 | 54 | 4.66* |
| L inferior parietal cortex |  | -42 | -46 | 54 |  |

**Table 3b.** Brain regions showing signification responses during the response phase of trials within the subgroup of participants that all chose (i.e. preferred) the drink labeled low-calorie at the end of the experiment (n=23; threshold: P < .001, k>10; *P_FWE_ < 0.05 at the whole-brain corrected cluster level).

| **Contrast/Region** | **Cluster**<br>**size**<br>**(voxels)** | **x** | **y** | **z** | **peak t** |
| --- | --- | --- | --- | --- | --- |
| ***Approach > avoid: preferred (low-calorie) > non-preferred (high-calorie) in subgroup that chose low-calorie drink (n=23)*** |  |  |  |  |  |
| L mid cingulate cortex | 62 | -12 | -30 | 46 | 6.03 |
| L postcentral gyrus |  | -8 | -30 | 54 |  |
| L postcentral gyrus | 51 | -40 | -16 | 40 | 4.88 |
| L inf parietal cortex | 17 | -42 | -28 | 40 | 4.74 |
| L sup frontal cortex | 22 | -14 | 42 | 30 | 4.63 |
| R cerebellum | 29 | 28 | -42 | -32 | 4.38 |
| L postcentral gyrus | 41 | -22 | -46 | 60 | 4.24 |
| L supramarginal gyrus | 18 | -58 | -26 | 36 | 3.8 |
| ***Approach > avoid: non-preferred (high-calorie) > preferred (low-calorie) in subgroup that chose low-calorie drink (n=23)*** |  |  |  |  |  |
| L inf/mid occipital cortex | 594 | -30 | -80 | -4 | 5.49* |
| R sup/mid occipital cortex | 72 | 16 | -86 | 20 | 4.88 |
| L dlPFC | 47 | 34 | 28 | 28 | 4.73 |
| L fusiform gyrus | 28 | -44 | -60 | -20 | 4.73 |
| L sup occipital cortex | 40 | -16 | -82 | 14 | 4.39 |
| L mid occipital cortex | 10 | -28 | -84 | 20 | 4.1 |
| R lingual gyrus | 33 | 14 | -84 | 4 | 4.06 |

In sum, our subjects demonstrated a greater approach bias behaviorally when anticipating the preferred (low-calorie) versus non-preferred (high-calorie) labeled drink, which was also chosen in more than 75% of the time when given the option between the two differently labeled (but identical) drinks after the experiment. At the neural level, the written information led to anticipation differences in a set of regions including the insula. During motivated action, the preferred drink elicited greater signal in the sensorimotor cortex during approach versus avoid actions relative to the non-preferred drink.

## Discussion

The present fMRI study examined the effects of differently labeling identical drinks on neural responses during drink anticipation and on behavioral and neural measures of motivated action – when participants had to make label-independent responses to obtain these drinks. We found that induced beliefs can exert influence on approach bias in brain and behavior, by labeling identical drinks as either ‘low-calorie’ or ‘high-calorie’. Although the liking scores did not differ between the labeled drinks, the vast majority of participants chose the drink labeled ‘low-calorie’ when given the option, which we interpret as a general preference for this label. The behavioral results are in line with this interpretation, as we observed an approach bias when anticipating the preferred drink (labeled as low-calorie) in both behavioral accuracy and reaction times. This interpretation was substantiated in a subgroup analysis in the participants that all chose the low-calorie drink. Our results are in line with a previous study that used similar joystick instructions (towards or away from the body), which showed automatic tendencies to approach appetitive foods (i.e. fruit) pictures and to avoid aversive pictures (i.e. rotten fruit; Piqueras-Fiszman et al., 2014). However, in our task, as in real life, approach behaviors towards food could result in more or less favorable outcomes. Our results extend the findings from a study in children, demonstrating that explicit motivation (i.e. grip force) was related to preferred food packaging of objectively identical products (Enax et al., 2015). Here, we show effects of labeling in an implicit, and perhaps more ecologically valid, way in adults, along with underlying neural mechanisms.

The absence of main behavioral effects of label indicates that the preferred drink did not become more salient overall, nor did participants generally perform better (i.e. became more goal-directed) for the drink they preferred. Instead, the labels affected the magnitude of the approach bias. We interpret our results in terms of approach bias because evaluative meaning attributed to the instructed joystick movement, namely ‘pull towards yourself’ and ‘push away from yourself’, is associated with approach and avoidance movements, respectively (Eder and Rothermund, 2008; Neumann and Strack, 2000). It has been suggested that both instrumental (e.g. habitual) and Pavlovian mechanisms contribute to the approach bias effect (Watson et al., 2012). For instance, cues with conditioned (Pavlovian) value can affect the vigor of instrumentally trained responses; an effect that emerges in the absence of a formal association between Pavlovian and instrumental contingencies (Talmi et al., 2008). In our instrumental version of the task, approach bias effects for the preferred, low-calorie labeled drink (i.e. errors for avoid actions) reflect an inability to inhibit an approach response, despite being rewarded with the actual preferred stimulus for a correct avoid response. Thus, our data show that participants were more motivated to approach the preferred stimulus and to avoid the less preferred stimulus, despite these automatic tendencies directly counteracting someone’s ability to obtain the associated, preferred reward. Top-down information could thus affect goal-directed control in hungry participants. Whether this label-specific decrease in goal-directed control is driven mostly by Pavlovian effects or by habitual, outcome-independent, instrumental responses remains unclear and could for example be investigated by using this instrumental version of the task before and after an outcome devaluation manipulation. Future research should also assess how our instrumental (and perhaps more ecologically valid) task relates to earlier findings of (non-instrumental) approach bias effects to food cues as a function of individual differences in eating behavior, e.g. health interest (Brignell et al., 2009; Havermans et al., 2011; van Rijn et al., 2016; Veenstra and de Jong, 2010).

Parallel to the behavioral effects, a stronger neural approach bias was observed when working for the preferred (low-calorie labeled) drink relative to the non-preferred (high-calorie labeled) drink in postcentral gyrus and superior parietal cortex. Supporting this interpretation, a subgroup analysis using the same interaction contrast in the participants that all preferred the drink labeled ‘low-calorie’ also yielded neural responses in a range of sensorimotor regions, although these findings were not significant after a whole-brain correction for multiple comparisons (pFWE<.05). This is probably due to a loss of power in the smaller subgroup (n=23 vs. N=31). This neural counterpart in sensorimotor cortex of the error rate effect therefore seems to reflect the greater efficiency with which the approach actions are executed when working for the preferred label. The finding that appetitive cues, such as food cues, can increase one’s propensity to act prior to the moment at which an instrumental response is required can underlie this effect, by increasing motor system excitability (Chiu et al., 2014; Freeman et al., 2014; Gupta and Aron, 2011). An alternative explanation would be that sensorimotor cortex is involved in the exertion of avoid-related effort. A separate region, but overlapping the left postcentral gyrus, was more strongly involved in avoid-relative to approach actions made with the right hand, similar to findings of a recent approach bias study (Radke et al., 2016). Interpreted in this way, the interaction between label and action found in neighboring (non-overlapping) sensorimotor and parietal cortices would reflect additional motor resource recruitment to overcome automatic approach biases for the non-preferred drink (Chong et al., 2017). The importance of cue-(or label-)induced motor responses is highlighted by studies in which motor responses toward or away from specific food cues were trained, which suggest that the motor component of approach biases towards foods can affect choice behavior (Stice et al., 2016). Similar to our findings, these previous motor system excitability and behavioral training studies point towards a strong role of the motor system in approach tendencies towards food.

Mere manipulation by word labels did not only result in behavioral and neural differences during approach and avoidance responses, but also in neural differences during presentation of the labels preceding the response. Specifically, anticipation of the non-preferred high-calorie versus the preferred low-calorie labeled drinks activated several temporal regions and the right ventral anterior insula. The (ventral) anterior insula is well-known for processing food rewards (Sescousse et al., 2013; van der Laan et al., 2011) and (anticipating previously experienced) taste differences (Dalenberg et al., 2015; Woods et al., 2011), but it is also associated with experiencing aversive smells (Wicker et al., 2003), anticipation of aversive stimuli (Nitschke et al., 2006b) and (aversive) taste learning (Bermúdez-Rattoni, 2014). Indeed, meta-analyses have shown the anterior insula to be responsive to (anticipation of) outcomes with both positive and negative subjective values (Bartra et al., 2013; Knutson and Greer, 2008). To dissociate between the interpretations of anticipated taste differences and aversive anticipation, we performed a subgroup analysis in the participants that all chose the drink labeled ‘low-calorie’ at the end of the experiment. This analysis revealed clusters in left anterior temporal cortex and left medial temporal lobe regions, including hippocampus and amygdala. These regions are involved in anticipatory memory processes for aversive events (Mackiewicz et al., 2006; Mechias et al., 2010).

Therefore, it seems likely that, in this subgroup, the anticipatory activations for the high-calorie vs the low-calorie anticipation is driven by a relatively larger aversive anticipation. Given that the insula results do not seem to be driven by the subgroup of participants that chose the drink with the label ‘low-calorie’, we interpret the anticipatory insula responses across the whole group in terms of anticipated taste intensity (Woods et al., 2011). It is unclear why we did not find increased responses for the reverse contrast (preferred > non-preferred label) in reward anticipation regions, such as the ventral striatum (Knutson et al., 2001). Perhaps the contrast was not strong enough to show this effect because of the identical objective, e.g. caloric, properties of the drinks. Indeed, the ventral striatum has been found to represent caloric values of foods independent of preference, whereas the insula was sensitive to differences in subjective properties (de Araujo et al., 2013).

Previous research on the neural basis of instructed beliefs point strongly towards a role for the prefrontal cortex in interpreting evidence in line with existing beliefs (Small, 2010) – across learning, decision and valuation processes (Biele et al., 2011; Doll et al., 2011; 2009; Engelmann et al., 2009; Li et al., 2011; Plassmann et al., 2008). We failed to observe prefrontal involvement in this study, which might be explained by the different task used here. In contrast to many of these previous studies, here we contrasted differently labeled drinks instead of contrasting the ‘actual’ with the believed truth, as one label was not more correct in relation to the actual sensory experience than the other. Also, here the actual sensory evidence was obtained during only 7 reward receipts following each block (the small number precludes analysis) instead of the many trials in the cited studies. Finally, participants in many of the previous studies had time for some deliberation on their responses, whereas the adaptive response deadline used here forces people to make immediate responses. Therefore, we might have tapped into faster, implicit processes rather than more deliberate control processes by the prefrontal cortex.

We observed an approach bias when anticipating a preferred, low-calorie label in both behavioral accuracy and reaction times and in sensorimotor cortex, despite the fact that our joystick task differed from most other approach-avoidance tasks in two important ways. First, as mentioned above, automatic tendencies to approach anticipated desired food stimuli directly counteracted someone’s motivation to obtain them in our instrumental task. Second, our task contained an (incentive) delay period between the stimulus to which a bias might exists and the moment of response, whereas usually the stimulus is shown at the time the response needs to be made (Rinck and Becker, 2007; Watson et al., 2012).

Nevertheless, our task was sensitive enough to detect differences caused by the food labels, showing that written information can influence task performance at a later point in time and despite contradicting participant’s (goal-directed) motivations. These findings suggest that approach-avoidance paradigms can be robust to time delays between cue and response and that future studies with this paradigm can attain increased ecological validity, by applying it in the motivational context of action-dependent outcomes.

The effect of misleading information on motivated action was consistent in brain and behavior. Conversely, responses after the neutral label resulted in a different pattern in brain and behavior. In contrast to the caloric drinks, the neutral drink was a different drink (water) that did not differ from the other drinks in subjective liking or wanting, making comparisons to the neutral label hard to interpret in terms of mere beliefs versus sensory or reward differences.

There is strong evidence that labels can influence food choices or subjective valuation, like willingness to pay, in experimental setups (Chandon and Wansink, 2012; Enax and Weber, 2015; Olson and Dover, 1978; Piqueras-Fiszman and Spence, 2015). However, in daily life, food choice might be more driven by implicit biases than in experimental environments. Here, we demonstrate that labels can indeed affect such implicit biases. Future research should investigate whether our behavioral and neural label effects are accompanied by altered food choice in daily life, in a wider range of products and labels.

## Conclusion

We report that written information in the form of food labels can lead to different approach tendencies behaviorally as well as in sensorimotor cortex for the preferred labeled drink. The subtle manipulation of written information led to approach bias effects that were strong enough to override goal-directed, instrumental responses to obtain the reward outcomes. These findings enhance our understanding of how expectancies caused by food labels affect motivational processes.

## Acknowledgements

This work is part of the FOCOM project, which was supported by the European Regional Development Fund and the Dutch Provinces Gelderland and Overijssel (Grant 2011-017004). We express our thanks to all partners of the FOCOM project for their valuable input. J.W. was also supported by a Food, Cognition and Behaviour grant of the Netherlands Organization for Scientific Research (NWO, grant 057-14-001). P.S. was supported by the European Union Seventh Framework Programme (FP7/2007–2013, Grant 607310). R.C. was supported by the James S. McDonnell Foundation (220020328) and a VICI grant (016.150.064) of NWO. E.A. was supported by a VENI grant of NWO (016.135.023).

